# Robotics-assisted acoustic surveys could deliver reliable, landscape-level biodiversity insights

**DOI:** 10.1101/2025.06.02.657416

**Authors:** Peggy A. Bevan, Cristina Banks-Leite, Mirko Kovac, Jenna Lawson, Lorenzo Picinali, Sarab S. Sethi

## Abstract

Terrestrial remote sensing approaches, such as acoustic monitoring, deliver finely resolved and reliable biodiversity data. However, the scalability of surveys is often limited by the effort, time and cost needed to deploy, maintain and retrieve sensors. Autonomous unmanned aerial vehicles (UAVs, or drones) are emerging as a promising tool for fully autonomous data collection, but there is considerable scope for their further use in ecology. In this study, we explored whether a novel approach to UAV-based acoustic monitoring could detect biodiversity patterns across a varied tropical landscape in Costa Rica. We simulated surveys of UAVs employing intermittent locomotion-based sampling strategies on an existing dataset of 26,411 hours of audio recorded from 341 static sites, with automated detections of 19 bird species (n = 1,819) and spider monkey (n = 2,977) vocalisations. We varied the number of samplers deployed in a single survey (sampling intensity) and whether the samplers move between sites randomly, in a pre-determined route to minimise travel time, or by adaptively responding to real-time detections (sampling strategy) and measured the impact on downstream ecological analyses. We found that observed avian species richness and spider monkey occupancy was not impacted by sampling strategy, but that sampling intensity had a strong influence on downstream metrics. Whilst our simulated UAV surveys were effective in capturing broad biodiversity trends, such as spider monkey occupancy and avian habitat associations, they were less suited for exhaustive species inventories, with rare species often missed at low sampling intensities. As autonomous UAV systems and acoustic AI analyses become more reliable and accessible, our study shows that combining these technologies could deliver valuable biodiversity data at scale.

## 1. Introduction

Terrestrial remote sensing approaches provide numerous advantages for biodiversity monitoring over traditional surveys, offering a non-invasive and scalable way to study ecosystems (Allan et al., 2018; Berger-Tal & Lahoz-Monfort, 2018; Lahoz-Monfort & Magrath, 2021; Stephenson, 2020). Among these, acoustic monitoring has proven particularly effective for surveying vocal species and quantifying human activity (Gibb et al., 2018; Ross et al., 2023) and has demonstrated great potential to contribute to biodiversity monitoring and policy compliance efforts (Stowell & Sueur, 2020; Sugai et al., 2019). Acoustic data can also be analysed rapidly using machine learning, either post-hoc (Lawson et al., 2023; Sethi et al., 2024) or in real-time (Sethi et al., 2018; Wägele et al., 2022). However, deploying acoustic sensors is a time-consuming and costly process, and survey locations can be limited by accessibility or safety concerns (Sethi et al., 2022).

Unmanned aerial vehicles (UAVs), or drones, are emerging as a transformative technology for ecological research (Robinson et al., 2022). For acoustic monitoring, recording data from sensors mounted on drones has the potential to improve scalability and reduce the logistical constraints associated with manually deploying static sensors. Early studies have explored UAVs equipped with microphones to record continuously during flight, using custom classification algorithms to detect bird sounds against the backdrop of engine noise (Michez et al., 2021; Wang et al., 2023; Wilson et al., 2017). However, the noise generated by drone motors can disturb wildlife (Wilson et al., 2017), and training novel detection algorithms demands more data, effort, and expertise than using established vocalisation detection models (e.g. BirdNET; Kahl et al., 2021). An alternative sampling approach is to land the UAV during recording periods and move between sampling sites on a defined schedule. This intermittent mobile sampling approach could mitigate noise-related disturbances, and more closely reflects survey designs of passive acoustic monitoring surveys, where static sensors are preferred over transect surveys for estimating animal activity or density (Browning et al., 2017; Lucas et al., 2015; Marques et al., 2013; Newson et al., 2017).

Most examples of UAV-based acoustic surveys have used UAVs which are manually controlled by trained pilots during missions. However, autonomous UAVs, which navigate themselves using sensors to avoid obstacles, could reduce barriers to access and enable finer resolution and larger scale biodiversity data collection. Lawson et al. (2024) designed an autonomous aquatic vehicle that could record sound above and below the water, collecting ecological data to the same standard as manual data collection methods. For terrestrial surveys, early stage prototypes have been developed to overcome challenges to automated acoustic surveys, such as navigating and landing in complex environments (Lan et al., 2024; Romanello et al., 2024; Zhou et al., 2022). Utilising these landing methods could reduce disturbance of wildlife and the environment by avoiding the need for any people to enter the survey area. However, it remains to be tested if deploying fleets of autonomous UAVs in the wild can deliver reliable ecological data, when compared to data from manually deployed sensors.

Furthermore, the deployment of autonomous UAVs for ecological surveys represents an opportunity to test novel survey designs that would not be possible with manual sensor deployment. UAV systems can be programmed with vehicle routing algorithms which optimise for a specific parameter, for example reducing the number of re-charges required (Guerber et al., 2021; Gunal, 2019; Macrina et al., 2020; Phalapanyakoon & Siripongwutikorn, 2021). Additionally, adaptive systems can respond to real-time input from sensors (Dwivedi et al., 2023; Hwang et al., 2019), a method successfully employed by underwater autonomous vehicles in tasks such as mapping the extent of underwater oil spills or locating phytoplankton blooms (Das et al., 2015; Jakuba et al., 2011). In biodiversity monitoring, adaptive sampling strategies have been used to detect environmental gradients or improve occupancy model accuracy (Flint et al., 2024; Henrys et al., 2024; Pacifici et al., 2016), but these occur over several survey periods, and do not respond to real-time animal detections. There is little understanding of how real-time adaptive sampling might impact the structure of ecological data, and if this could produce robust measures of biodiversity compared to more established survey techniques.

Designing a UAV-based acoustic survey presents unique limitations, particularly regarding sampling completeness. Standard passive acoustic monitoring surveys typically deploy one recording device at each sampling site, and all devices record data simultaneously over extended periods. In contrast, intermittent sampling by UAVs means that the recording duration at any given site is restricted. As some animal vocalisations are temporally dependent, sampling at the same time of day across independent sites is important for robust comparisons of species detections across sites. It remains unclear to what extent incomplete sampling and reduced sampling evenness might affect estimates of species richness, community composition, or habitat associations. Addressing these gaps is critical to understanding the trade-offs associated with UAV-based acoustic monitoring and to help practitioners choose the optimal sampling strategy to meet their survey aims.

In this study, we ask how reduced effort resulting from an intermittent sampling protocol would impact downstream biodiversity metrics. We designed and ran simulated UAV surveys using an existing acoustic dataset from the Osa Peninsula, Costa Rica. This dataset has previously been analysed to measure avian (Sethi et al., 2024) and spider monkey (Lawson et al., 2023) diversity and occupancy, respectively. Using published data from these studies, we simulated surveys under four sampling strategies that could be carried out by a network of autonomous UAVs—random, routed, and two types of adaptive sampling. We evaluated the performance of each strategy under varying levels of sampling intensity, defined by the numbers of UAVs (samplers) deployed in a mission. By replicating key analyses from the original studies, we assessed the impact of intermittent sampling on estimates of avian species richness and spider monkey occupancy over a forest cover gradient. This work is the first to consider the full pipeline of UAV-based acoustic surveys and to investigate the impact of sampling techniques on downstream ecological metrics, informing future applications of this technology in ecological research.

## 2. Materials & Methods

### Study area & sampling design

For our simulations, we used species vocalisation detections from two existing datasets from a large passive acoustic monitoring survey in the Osa Peninsula, Costa Rica (Lawson et al., 2023; Sethi et al., 2024). The Osa Peninsula, located on the south pacific coast of Costa Rica, contains biodiverse tropical broadleaf evergreen lowland rainforest (Arturo Sánchez-Azofeifa et al., 2003; Gilbert et al., 2016), embedded within a mosaic of pasture, plantations and urban centres (Fig. 1).

**Fig. 1.**
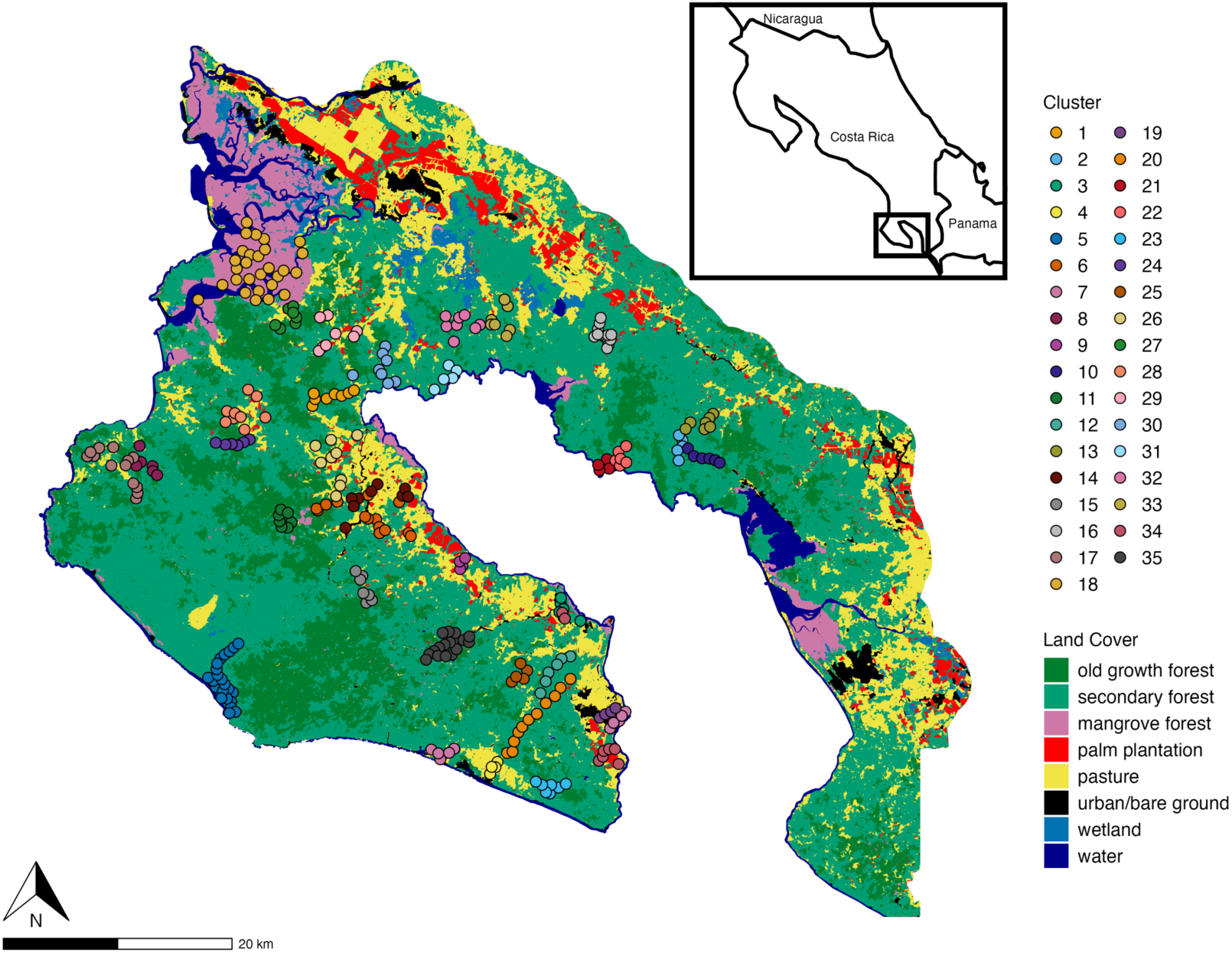
Map of Survey Area in the Osa peninsula, Costa Rica. Each coloured point represents a sampling location where a single acoustic recorder was placed. Acoustic recorders were deployed in ‘clusters’ across the landscape, and points are coloured accordingly. This is also the sampling unit used for simulated drone-based surveys. The Land Use Map shows the land use categories in the region, created at a scale of 5 × 5m using Landsat 5 Thematic Mapper (TM) and Landsat 8 Operational Land Imager (OLI) (Shrestha et al., 2018).

We chose this dataset for our drone simulations because of the sampling design. Due to access limitations, a uniform sampling design across the survey area (1093 km^2^) was not possible. Instead, a set of sampling groups were created which appear as ‘clusters’ (Fig. 1). This mimics the sampling design of a robotic UAV survey, where samplers begin from central charging station, and travel to recording locations within a set radius of the charging station. The dataset contained recordings from 341 sites grouped into 35 clusters, with an average of 8.7 sampling sites within each cluster (range [4, 28]).

Sampling sites were stratified across five land-use categories, with the number of recorders placed in each land-use category representative of its percentage cover across the region (determined from the 5 x 5 m Landsat 5 Thematic Mapper (TM) and Landsat 8 Operational Land Imager (OLI) (Lawson et al., 2023; Shrestha et al., 2018). The land-use categories were old growth forest, secondary forest, palm plantation, teak plantation and grassland. At each sampling cluster, the first recorder was placed by walking 500m in a random direction, and remaining recorders placed at a minimum of 500m apart to maximise independence between samples (Figueira et al., 2015). Where possible, trails were not used to avoid bias, however, where this was not possible, devices were placed a minimum distance of 200m perpendicular to a trail. Recording devices were also placed at a minimum distance of 200m from habitat boundaries, to ensure sounds were solely from the classified habitat. Recordings were obtained using AudioMoth devices (Open Acoustics Devices, UK). Recorders ran for a minimum of seven consecutive days (range [7, 16 days]) to allow for variability in activity across different days and to allow for sufficient sampling effort. The devices were set to record on a schedule of 05:00–09:30, 14:00–18:30 and 21:00–03:00, a total of 15 h each day. Data was recorded constantly during these periods at a sample rate of 48 kHz (Bradfer-Lawrence et al., 2019). Sampling was conducted during the dry season (December-August). The final dataset included 26411 hours of uncompressed 16-bit audio files.

### Data processing

#### Avian species richness

We used avian vocalization detections from Sethi et al. (2024), who used an open-source bird vocalisation detection model, BirdNET (Kahl et al., 2021), to identify bird calls in the audio data. BirdNET is a convolutional neural network (CNN) trained on bird vocalizations from many online call libraries. The BirdNET application clips audio in to three second chunks and converts to a mel-scale spectrogram before passing through the CNN to give a list of predicted species detections, with a prediction probability. Sethi et al. (2024) had the geographic species filter enabled and set a model confidence threshold of 0.8. For this study, we kept detections for 19 species which the model was able to detect with > 90% precision (*i.e.* low false positive rate) based on verification of a random subset of 50 detections by an experienced local ornithologist (Sethi et al., 2024). This resulted in 1819 bird detections across 126 sites and 32 clusters. The 19 species represents a very small proportion of the ∼465 bird species that can be found on the Osa peninsula, and is a result of the geographical and taxonomic bias in the training datasets of BirdNET (Stowell, 2022). It is important to note that analyses of species richness from this study are not intended to make conclusions about avian distribution in the Osa Peninsula, but to be used as an example of real species spatiotemporal patterns and dynamics.

#### Spider monkey occupancy

We used spider monkey detections from Lawson et al. (2023), who trained a deep learning convolutional neural network to detect the species’ vocalisations. Lawson et al. (2023) manually annotated 561 examples from 13 sites of the ‘whinny’ vocalisation made my spider monkeys. The neural network used, proposed by Rizos et al. (2021), used a deep, convolutional neural network architecture. All spider monkey detections were manually verified to ensure all false positives were removed. The final dataset contained 2977 true positives across 64 of 341 sites (Lawson et al., 2023). These detections were converted into a 7-day detection history per site with each day coded as 1 or 0 to represent presence or absence of spider monkeys. Data at a finer temporal resolution was not published, limiting the scope of our simulations for the spider monkey dataset.

#### UAV survey simulations

To evaluate data collectable by an autonomous UAV system, we ran a series of simulations to collect data from the bird and spider monkey vocalisation detection datasets. Surveys were designed as intermittent mobile sampling, meaning that data collection (*i.e.* sound recording) occurs when the drone is stationary at a site to minimise the impact of flying noise on recording quality and animal disturbance.

Such intermittent locomotion can be achieved by landing on the ground or perching in the canopy (e.g. Zheng et al., 2023). All simulations were run in Python 3.

UAV functionality was based on the DJI Mini 2, a commercially available multi-rotor drone. This device can achieve approximately 30 minutes of flight time when unweighted. As UAVs would be weighted with a microphone and related hardware (approx. 80g including separate batteries) (Hill et al., 2019), we estimated a maximum flight time of 20 minutes (1200 seconds). Battery consumption was calculated as a function of flight time. Average flying speed was set as 15 meters/second, and an extra penalty of 16 seconds was added per take-off to account for the increased battery consumption of increasing the UAV’s altitude. We assumed that while stationary and landed the UAV consumed no battery. We did not account for the impact of wind, temperature and other factors which may reduce overall battery in a real-life scenario since we did not have this data at appropriate resolutions.

Simulations were run at the level of a sampling cluster (Fig. 1). The sampling site closest to the mean latitude and mean longitude of all sampling sites in the cluster was selected to be a ‘charging station’. Each simulation began with all samplers (an autonomous UAV equipped with microphone) at the charging site in the first timestep. At each sequential timestep, samplers would be moved to other sampling locations within the cluster, and ‘collect’ any detections from the real audio dataset at that timestep and location. A timestep was 1 hour for the avian dataset simulations (15 timesteps per day, ranging 7-16 days) and 1 day for the spider monkey dataset simulations (7 timesteps in total). If a sampler did not have enough battery to both visit the next selected site and then return to the charging station, the sampler was sent directly back to the charging station. All simulations were repeated 50 times to account for stochasticity in site selection.

Our simulations experimented with two parameters: *sampling intensity* and *sampling strategy*. *Sampling Intensity* represents the number of samplers deployed simultaneously. We used five values of sampling intensity (0.2, 0.4, 0.6, 0.8, 1), where each value represents number of samplers as a proportion of the total sites in the cluster. A sampling intensity value of 1, representing complete sampling (*i.e.* one sampler per sampling site), was used for baseline comparisons.

*Sampling strategy* defined how the samplers’ route between sites was determined. We created four strategies: *Random, Routed, Adaptive explorative and Adaptive exploitative*.

In the random sampling strategy, the choice of next site was determined by random selection with replacement, allowing the same site to be chosen in consecutive timesteps. Sites already selected by another sampler were removed from the list of available sites, to avoid multiple visits to the same site at the same time.

For the routed sampling strategy, sampling sites at each cluster were grouped into geographically distinct sub-groups using *k*-means clustering based on their GPS coordinates, where *k* = *N_samplers_*. Each sampler would only travel to sites within a specific sub-group. At each timestep, the choice of next site was based on the number of previous visits, prioritizing sites that had been sampled the least. Among these sites, the nearest available site was selected. This strategy meant that over the course of the survey, all sites in a cluster were visited equally, and the order of site visits remained relatively consistent because the nearest-neighbour algorithm repeatedly identified the same shortest route between sites.

The third and fourth sampling strategies, adaptive explorative and adaptive exploitative, considered a scenario where samplers had on-board processing capability such that animal vocalisations could be detected in real-time, and adapt the choice of next site. This strategy was run for the avian community dataset only, due to the higher temporal resolution of this dataset. Adaptive sampling can be configured in many ways, depending on the metric being optimised for *e.g.* finding a gradient (Hwang et al., 2019) or increasing occupancy confidence (Pacifici et al., 2016). Here, we chose to optimise sampling effort to increase confidence that all species present in each cluster were detected. To achieve this, we programmed the routing system to prioritise sites where more frequent avian vocalisation rates were detected, as these were expected to represent areas of high avian activity. For the first two days of the simulation (30 timesteps), sites were sampled using the routed sampling strategy explained above, to ensure baseline coverage of all sites. After 30 timesteps, the choice of next site was decided through random selection with replacement, where the probability of choosing each site was biased by the number of vocalisations previously detected and total number of visits to that site. The probability weighting of site *i* (*P*(*i*)) was determined as:

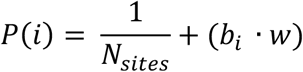

Where the bias, *b_i_* is calculated as total number of vocalisations previously detected at site *i*, divided by the total number of visits to site *i*, normalised by the sum of these values for all available sites. *w* represents a weighting factor that determines the disparity between high and low probabilities.

We tested two values of *w* to explore the difference between an ‘explorative’ adaptive system (*w* = 0.3), where real-time detections give a low bias to site choices, and an ‘exploitative’ adaptive system (*w* = 100), where real-time detections give a high bias to choice of next site.

#### Evaluating simulation performance

Simulation evaluation was conducted in R (V4.4.1). For simulations using the dataset of BirdNET detections, we measured the total number of species detected (species richness) across the survey area and in each land-use type (Old Growth Forest, Secondary Forest, Mangrove Forest, Grassland, Palm Plantation, Teak plantation) and compared these values to those collected through complete sampling (sampling intensity = 1). We used a generalised linear mixed-effects model (GLMM) with a Poisson distribution to determine the effect of sampling strategy (random, routed, adaptive explorative and adaptive exploitative) and sampling intensity on species richness. Iteration number was included as a random effect.

For the spider monkey dataset, we used the 7-day detection history to run a logistic GLM that predicted probability of spider monkey presence in response to forest cover, replicating the analysis from Lawson et al. (2023). To define a minimum threshold of forest cover for spider monkey occupancy, we used the receiver operating characteristic (ROC) curve to determine a cut-off point that maximised specificity and sensitivity for predicting positive occupancy. We then predicted occupancy probability over the range of forest cover, and found the value of forest cover that intersected with the cut-off probability to get the minimum forest cover threshold. This threshold is not the minimum value spider monkeys were detected at, but more closely represents the value of forest cover above which spider monkeys are consistently detected, therefore accounting for anomalous detections in areas of lower forest cover. We calculated the minimum forest cover threshold for each simulation and compared to values from complete sampling (sampling intensity = 1).

#### Survey logistics

To show how changing sampling strategy and intensity would impact other aspects of a UAV-based survey design, we measured sampling evenness, total sampler travel time and the number of recharges for all simulations on the avian vocalisation dataset.

To measure sampling evenness across space and time, we calculated the frequency of visits to each site at each hour in the day the survey was active (05:00–09:30, 14:00–18:30 and 21:00–03:00, n = 15 hours) and calculated Shannon evenness index (SEI) for each cluster. The SEI value is bound between 0 and 1, where a higher value indicates more equal visits to a site across hours in the day. We calculated SEI for each cluster and averaged this across all 35 clusters for each value of sampling strategy and sampling intensity.

Average total travel time was calculated by summing the total distance travelled by all samplers in a cluster during a single simulation then normalised by the number of sites, so that travel time was comparable across clusters with different numbers of sites. The same method was used to record the average number of visits to the charging station.

## 3. Results

### Simulation results: Routed and Random Sampling Strategies

All simulations were able to recover the patterns of BirdNET species detections across land-use types, although species richness reduced with sampling intensity (Fig. 2). Across all values of sampling intensity, the measured species richness was always highest in grassland, followed by secondary forest, old growth forest, palm plantation, mangroves and teak plantation (Fig. 2A). Sampling strategy (random or routed) had no impact on measured species richness.

**Fig. 2.**
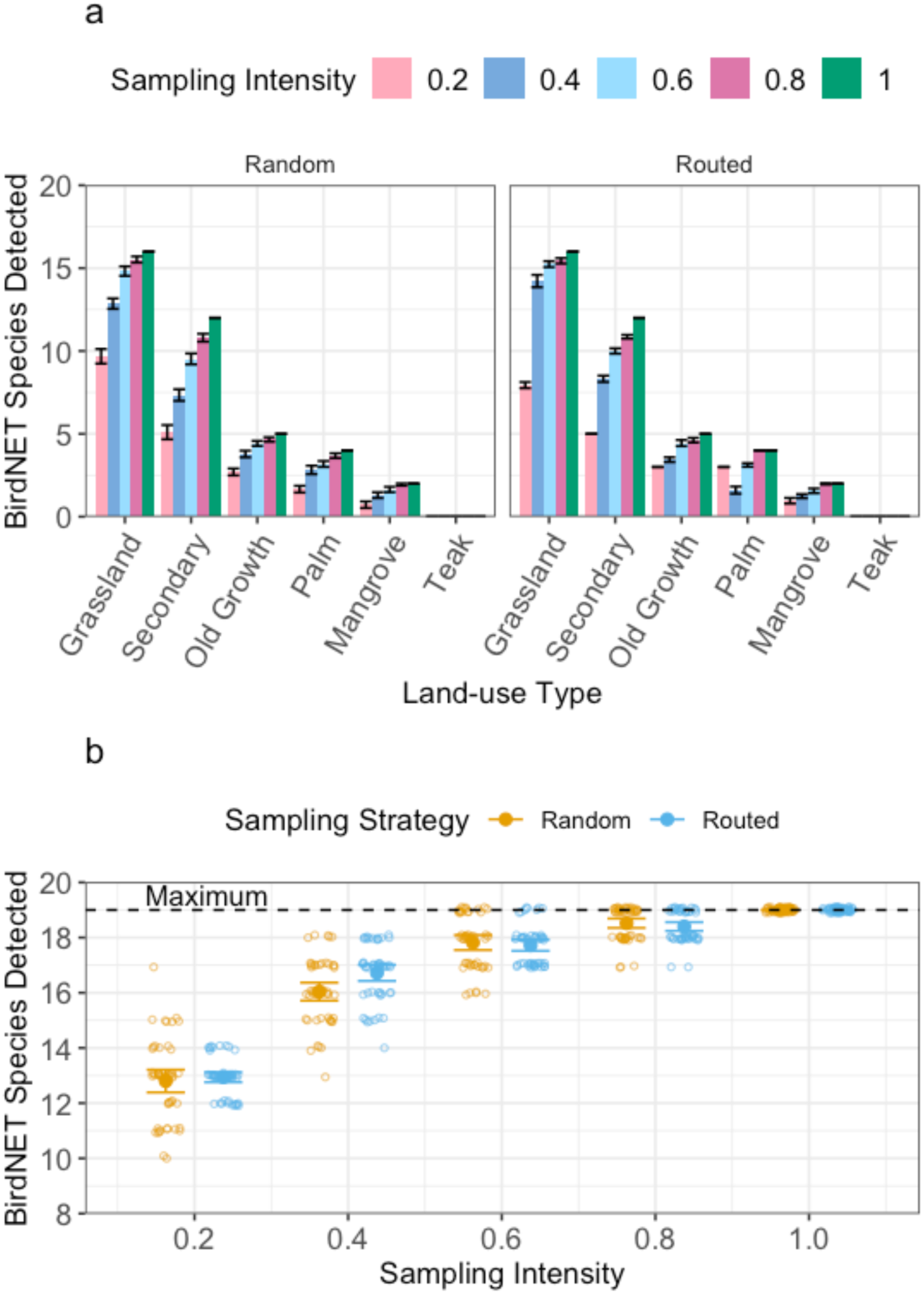
Patterns of biodiversity across land-use types detected with low number of UAV samplers, but uncommon species hard to detect. Total BirdNET detections from UAV survey simulations that varied the number of samplers deployed simulataneously (sampling intensity) and the sampling strategy (random movement between sites, or Routed, optimising for the shortest travel distance). a) the number of species detected by BirdNET across five land-use types. b) the total number of species detected by BirdNET across the entire survey (maximum 19). Note that species numbers and habitat use patterns are not indicative of real biodiversity patterns in the study area.

When examining the full list of BirdNET species detections across all sites, no simulations with a sampling intensity <1 were able to consistently detect the maximum species richness (n = 19 species), although some iterations at 0.6 and 0.8 sampling intensity did detect the maximum species richness (Fig. 2B). Results of the GLM showed no impact from sampling strategy on measured species richness (*p =* 0.58) but sampling intensity had a strong positive effect (*p* < 0.001).

For the spider monkey simulations, we found that the minimum forest cover threshold for predicting spider monkey occupancy tended to be overestimated with high uncertainty, although this improved with sampling intensity. The results from complete sampling (sampling intensity = 1) predicted a minimum threshold of 0.909, equivalent to 91% forest cover. This value was not consistently achieved by any of the simulations, regardless of sampling strategy. For simulations with the lowest sampling intensity (0.2) and random sampling strategy, the average forest cover threshold was 95%, ranging between 91% and 100% (Fig. 3). Interestingly, the raw minimum level of forest cover that a spider monkey was detected at (79%) was detected by both sampling strategies at all sampling intensities except for 0.2.

**Fig. 3.**
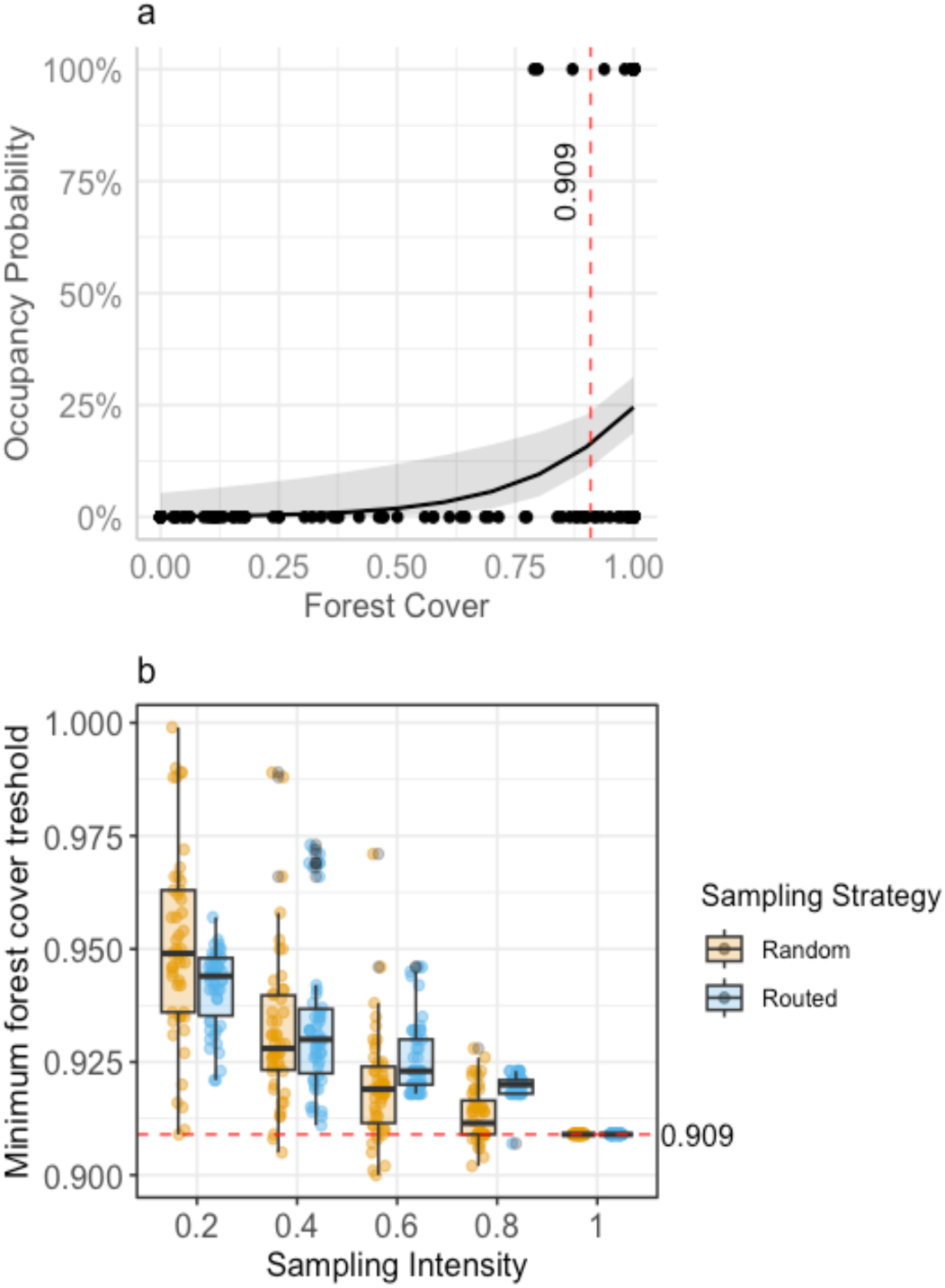
UAV-based acoustic surveys overestimate the minimum forest cover requirements for spider monkeys. a) The predicted spider monkey occupancy over a gradient of forest cover within 100m of the sensor when the entire dataset was included (Sampling Intensity = 1), as presented in Lawson et al., (2024). The minimum forest cover threshold, calculated by finding the cutoff point that maximised the probability of a positive spider monkey detection, was 0.909 (red dashed line). b) Shows a repetition of this analysis using data collected from simulated UAV surveys, where sampling intensity (the number of drones deployed simultaneously, as a proportion of total sites) and sampling strategy (Random movement between sites or Routed movement that prioritises nearest neighbours) were varied. Boxplots summarise the minimum forest cover threshold estimated from 50 iterations of each simulation, where the box represents the lower quartile, median and upper quartile, and the whiskers are 1.5 * the Interquartile Range. Points show the raw results from each simulation.

### Simulation results: Adaptive Sampling

Including knowledge of real-time detections into the routing algorithm did not show an improvement on random sampling when looking at total number of species detected per survey. We investigated two adaptive routing strategies in our simulations by adapting the weighting parameter, *w*, in the routing algorithm: explorative (*w* = 0.3) and exploitative (*w* = 100). At very low sampling intensity (0.2), the adaptive exploitative strategy detected a higher species richness than the adaptive explorative strategy on average, but estimates of species richness were no higher than the random sampling strategy (Fig. 4). Results of the GLM showed that sampling intensity resulted in higher species richness (Est = 0.44, *p* < 0.001) but none of the tested sampling strategies had any effect (explorative: *p* = 0.58; exploitative, *p* = 0.85).

**Fig. 4.**
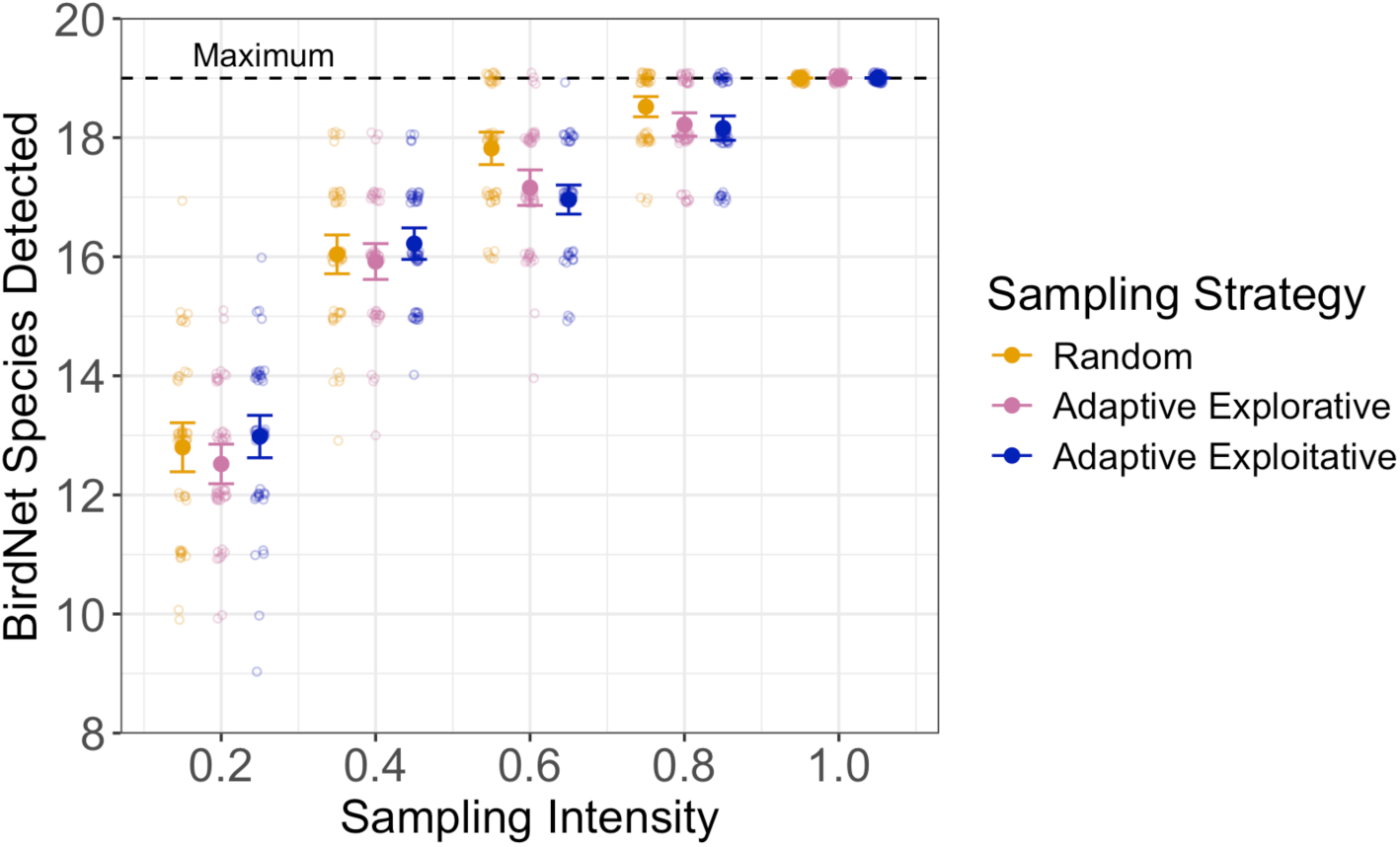
Incorporating very simple real time detections into route planning does not detect more species than a random sampling strategy. Unique BirdNET species detections from three simulations of an existing acoustic dataset. In each simulation, we varied sampling intensity (number of UAVs deployed simultaneously, as a proportion of total sites) and sampling strategy, defined as random movement between sites or two types of adaptive sampling, where samplers change routes depending on real-time detections.

Adaptive explorative indicates that previous vocalisation frequency at a site gives a low bias towards visiting that site again over others, whereas adaptive exploitative indicates vocalisation frequency at a site will give a high bias towards visiting that site again over others. Each solid point represents the average number of species detected over 50 simulation iterations, with error bars representing 95% confidence intervals. The faded points show the actual species richness from each iteration.

### Survey logistics

We found that sampling strategy had little impact on sampling evenness of a survey (how evenly samples were spread across hours in the day and sites per cluster over the full sampling period). We found no difference in sampling evenness between random, routed or adaptive sampling strategies (Fig. 5a). However, the variation in sampling evenness across the 50 simulation iterations was larger in the routed simulations than random or adaptive. Although increasing sampling intensity improved the average sampling evenness score, the lowest value of sampling evenness was 0.95. This means that even high manipulations to the sampling regime had a low impact on overall sampling evenness and would have little impact on the ability to straightforwardly compare sampled data across sites without adding complexity to downstream analyses.

**Fig. 5.**
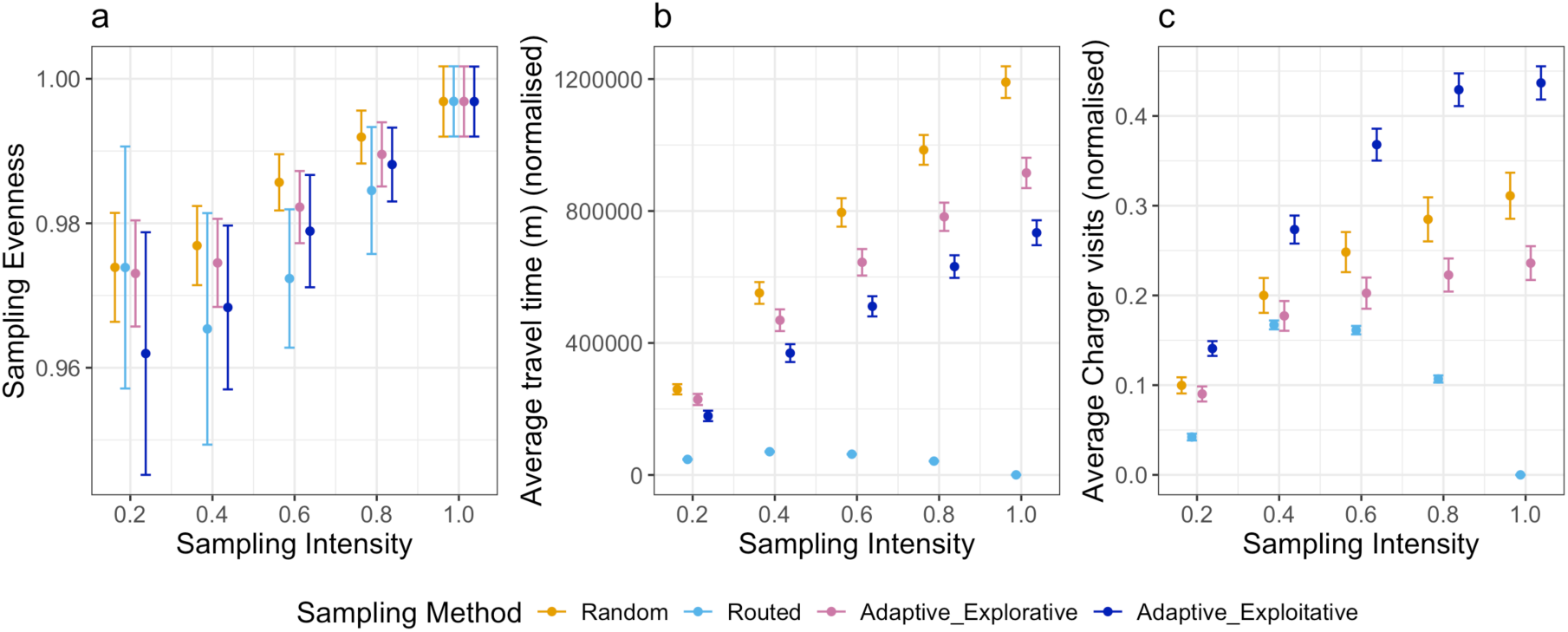
Distance based, routed sampling improves energy consumption but not sampling evenness. Comparison of survey logistics for different drone survey simulations where sampling intensity (number of UAVs deployed simultaneously, as a proportion of total sites) and sampling strategy, defined as random movement between sites, routed (optimising for the shortest travel distance) or two types of adaptive sampling, where samplers change routes depending on real-time detections. Adaptive explorative indicates that previous vocalisation frequency at a site gives a low bias towards visiting that site again over others, whereas adaptive exploitative indicates previous vocalisation frequency at a site will give a high bias towards visiting that site again over others. We calculated a) sampling evenness using Shannon’s evenness index (SEI), where a high value represents even sampling across sites and hours in the day. b) Average total travel time, normalised by total number of sites in a cluster and c) average number of charger visits, normalised by total number of sites in a cluster. For all plots, each point represents the average of 50 simulation iterations, with error bars representing 95% confidence intervals..

When looking at total energy consumption of a drone survey, using the routed sampling strategy reduced the average travel time per sampler (Fig. 5b) and total number of charger visits per sampler (Fig. 5c). This was in high contrast to the distances travelled by drones in both the random and adaptive sampling strategies. The difference in travel time and charger visits between routed strategies and the other strategies was higher at higher sampling intensities, likely due to the creation of sub-groups that forced samplers to stay in one geographical area.

## 4. Discussion

This study demonstrates that the intermittent sampling design of a UAV-based acoustic monitoring survey can effectively detect broad biodiversity patterns such as associations with land-use types, but is limited in its ability to capture complete species lists at low sampling intensities. The ability of a UAV-based acoustic survey to detect landscape-level biodiversity patterns indicates the potential of this approach as a preliminary survey tool for identifying sites of high conservation value. For example, UAVs could be used to rapidly assess biodiversity gradients across land-use types, reducing the need for manual deployments of acoustic sensors which could take a team of researchers days to weeks depending on the scale. Across all sampling strategies, we found that the primary factor that impacted species detection accuracy was the overall sampling intensity (number of UAVs deployed simultaneously) rather than the specific sampling strategy employed. Whilst previous studies on drone routing have largely focused on optimizing for efficiency in logistical applications (Macrina et al., 2020), this is the first study to investigate the impact of intermittent sampling strategies on the quality of downstream biodiversity data.

One key consideration of survey design is the evenness of sampling across space and time, to ensure both between- and within-site variability can be straightforwardly detected (Rhodes & Jonzén, 2011). Incomplete sampling across either of these axes can lead to bias in biodiversity metrics like species richness (Banks-Leite et al., 2012; Chave, 2013; Field et al., 2002; Lahoz-Monfort et al., 2014). We initially hypothesized that the routed sampling strategy, designed to distribute visits evenly across sites would enhance sampling evenness. However, we found that sampling strategy had minimal influence on sampling evenness. This suggests that even simple, non-optimized sampling strategies can achieve sufficient site coverage, provided sampling intensity is high enough. Thus, the results of our experiments suggest the ability of UAV-based surveys to provide representative biodiversity metrics is largely dependent on the number of samplers deployed or the length of total survey period, rather than the routing systems tested here.

Real-world deployments of UAV-surveys may be influenced by additional factors such as the behavioural response of animals to UAVs or weather conditions, that we were not able to control for. Previous studies on UAV disturbance have mainly attempted to determine an optimal flight height that minimises wildlife disturbance (Corcoran et al., 2021; Duporge et al., 2021; McEvoy et al., 2016). However, the survey design proposed here involves landing drones before recording data, so the recovery time of wildlife to short term disturbance is important to consider. Weston et al. (2020) found that a UAV take-off or landing will elicit escape responses in birds within 40m, but it has also been shown that bird vocalisation rate can recover in less than 4 minutes after a noise disturbance like a snowmobile (Cretois et al., 2024). The level of tolerance of a species to drone noise will also depend highly on the focal taxa and UAV flight path (Afridi et al., 2024; Mesquita et al., 2022). We recommend that UAV-based acoustic surveys incorporate a latency period after landing to minimize potential disturbance effects, but further research is needed to understand recovery times of wildlife to UAV landing.

Adaptive sampling is not yet widely used in biodiversity surveys, but advances in autonomous robotics present new opportunities to explore the potential of this approach (Henrys et al., 2024). The adaptive sampling strategy employed here showed no improvement in detecting species over randomly sampling sites. This could be linked to survey design. In our study, species detections were most frequent in grassland habitats. The sampling clusters that UAVs moved within were mostly made up of a single land-use type (Fig. 1). This meant that for many clusters, there was low variation in between-site species richness, and the adaptive sampling was not able to find sites with high species richness. This strategy may work better in more fragmented landscapes, where adaptive sampling could help to expose gradients in biodiversity. Furthermore, the adaptive strategy we employed was relatively simple, only accounting for one variable. Boubrima and Knightly (2021) created an adaptive UAV system that correlated air pollutant measures with wind, temperature and humidity to map the extent of pollution. A more advanced adaptive system for ecological surveys could incorporate real-time measures of abiotic (e.g. precipitation) or biotic (e.g. land classification) factors to influence route planning.

In conclusion, our findings highlight the feasibility of UAV-based acoustic monitoring as a scalable approach for biodiversity assessment. We demonstrate that the number of UAVs deployed simultaneously plays a greater role in determining detection success than the choice of sampling strategy, though routing will reduce power consumption. As UAV technology continues to advance, further research is needed to explore the applications of autonomous UAV-based ecological surveys across different ecosystems, taxonomic groups and data collection methods (*e.g.* camera trapping, eDNA). In addition, future studies should prioritize developing novel sensor placement methods and autonomy frameworks that are validated in field conditions to maximize their potential for conservation science.

## Acknowledgements

We thank the UK Acoustics Network Plus (UKAN+) for providing funding to complete this research. We would also like to thank the staff at El Ministerio de Ambiente y Energia (MINAE) and La Sistema Nacional de Áreas de Conservación (SINAC) for their support with permits and for the field support from their park rangers. This work was partially supported by funding from the Engineering and Physical Sciences Research Council, the SERI funded ERC ProteusDrone consolidator grant (grant no: MB22.00066) and the Empa-Imperial College London research partnership. Mirko Kovac was supported by the Royal Society Wolfson fellowship (RSWF/R1/18003). Jenna Lawson was supported by the Natural Environment Research Council (grant no. NE/L002515/1).

## Data availability

All our code is available on the following anonymised GitHub repository: https://anonymous.4open.science/r/PAM_drones_simulator-3633. All data used is publicly available at https://zenodo.org/records/6511837 (From Lawson et al., 2023) and https://zenodo.org/records/10820823 (Sethi et al. 2024).

